# Transcutaneous Vagus Nerve Stimulation Boosts Post-Error Accuracy During Perceptual Decision-Making

**DOI:** 10.1101/2024.10.21.619457

**Authors:** Shiyong Su, Thomas Vanvoorden, Pierre Le Denmat, Alexandre Zénon, Clara Braconnier, Julie Duque

## Abstract

The locus coeruleus-norepinephrine (LC-NE) system is a well-established regulator of behavior, yet its precise role remains unclear. Animal studies predominantly support a “gain” hypothesis, suggesting that the LC-NE system enhances sensory processing, while human studies have proposed an alternative “urgency” hypothesis, postulating that LC-NE primarily accelerates responses. To address this discrepancy, we administered transcutaneous vagus nerve stimulation (tVNS) in two experiments involving 43 participants. In the first experiment, we showed that 4-second tVNS trains reliably induced greater pupil dilation compared to SHAM condition, indicating increased LC-NE activity. In the second experiment, we applied tVNS during a random dot motion task to assess its impact on perceptual decision-making. Notably, tVNS improved accuracy without affecting reaction times, which appears inconsistent with the “urgency” hypothesis. Drift-diffusion model analyses further supported the “gain” hypothesis, revealing that tVNS increased the drift rate, indicative of enhanced evidence accumulation. Accuracy and drift-rate improvements were especially pronounced following errors and in less proficient participants, who otherwise exhibited post-error declines in these measures under SHAM condition. Our findings suggest that the influence of the LC-NE system adapts to task demands, becoming especially beneficial in challenging contexts. Overall, this study underscores the potential of tVNS as a non-invasive tool to investigate the causal role of the LC-NE system in human behavior.

## Introduction

The locus coeruleus (LC), a small brainstem nucleus, is the main source of norepinephrine (NE) in the brain, exerting widespread influence over cortical and subcortical brain areas. Traditionally, LC plays a central role in regulating arousal which is known to impact behavior [1-3]. Dysfunctions in LC-NE activity have been implicated in the etiology of attention-deficit/hyperactivity disorder (ADHD) [4], anxiety [5], depression [6], and the cognitive decline in Alzheimer’s and Parkinson’s diseases [7, 8]. The extensive projections of LC and its involvement in major neurological disorders emphasize the need to better understand its role in behavioral control.

Animal studies using various recording and interference methods have significantly advanced our understanding of LC-NE function [9-14]. Evidence suggests that LC-NE activity enhances the signal-to-noise ratio of sensory response across modalities, including visual [15, 16], olfactory [17], somatosensory [18, 19], and auditory signals ([19, 20]; see [21] for different findings). Interestingly, such fine-tuning of sensory response is notably beneficial for behavior, as heightened LC-NE activity has been associated with improved performance during perceptual decision-making [22, 23]. These findings have prompted a hypothesis that LC-NE activity enhances the “gain” of perceptual processing [24-26], with a probable spotlight on task-specific processes considering the brain’s finite energy resources [9, 27, 28]. However, it is important to note that excessive LC-NE activity can be detrimental, following an inverted U-shaped curve [29].

Human studies have also explored the role of the LC in behavior, often by tracking pupil size as an indicator of LC-NE activity [30]. This research spans various behavioral contexts, including perceptual decision-making [31-33]. One key finding is that pupil dilation increases when subjects emphasize fast (but then less accurate) decisions under time pressure compared to when they prioritize accuracy by slowing down [34]. This has led to an alternative “urgency” hypothesis [35], suggesting that LC activation creates urgency, prompting quicker decisions at the cost of accuracy, which contrasts with the “gain” hypothesis prevalent in animal research. However, some evidence does indicate a connection between pupil size and the efficiency of sensory processing in humans [36-38], making it unclear whether LC primarily enhances “gain” or creates “urgency” [39] during perceptual decision-making.

In this study, we aimed to clarify LC function in humans using transcutaneous vagus nerve stimulation (tVNS), a non-invasive method to increase LC-NE activity [40-43]. We conducted two experiments with 43 participants. The first experiment demonstrated the efficacy of a novel tVNS protocol in inducing short-term pupil dilation at rest, indicating a temporary increase in LC-NE activity. In the second experiment, we applied this protocol during a random dot motion (RDM) task to investigate the causal role of LC in perceptual decision-making. According to the “gain” hypothesis, increased LC-NE activity by tVNS should improve accuracy, while the “urgency” hypothesis predicts faster but less accurate decisions. These hypotheses are also distinct within the drift-diffusion model (DDM) of decision-making [44]. In the DDM, binary decisions are represented as the noisy accumulation of sensory evidence until one of two decision boundaries is reached. The “gain” hypothesis predicts that tVNS should increase the drift rate, representing more efficient evidence accumulation [45] [46], while the “urgency” hypothesis suggests tVNS would lower the initial boundary and/or accelerate boundary collapsing, indicating greater urgency [47]. Our findings in the RDM task support the “gain” hypothesis, showing a selective contribution of LC to accuracy, particularly when performance is likely to decline after errors.

## Materials and Method

### Participants and experimental setup

A total of 43 healthy subjects participated in two experiments conducted in a quiet room with moderate light. Participants sat in front of a computer screen, with their heads positioned on a chin rest (see details in Fig. 1A). Pupillometry was recorded throughout both experiments, and participants were instructed to minimize movements and blinks during the recording. In Experiment 1, 22 subjects (13 women, 24 ± 3.7 years) were tested to characterize the effect of a 4-second tVNS train on pupil size at rest. In Experiment 2, the remaining 21 subjects (12 women, 26 ± 4.5 years) underwent the same tVNS procedure during a RDM task. All participants had normal or corrected vision, no history of neurological or psychiatric disorders, and no physical injuries. They provided written informed consent and received compensation for their participation. The study was approved by the Ethics Committee of the Université catholique de Louvain (reference: 2023/13JUL/322) and adhered to the Declaration of Helsinki.

**Figure 1.**
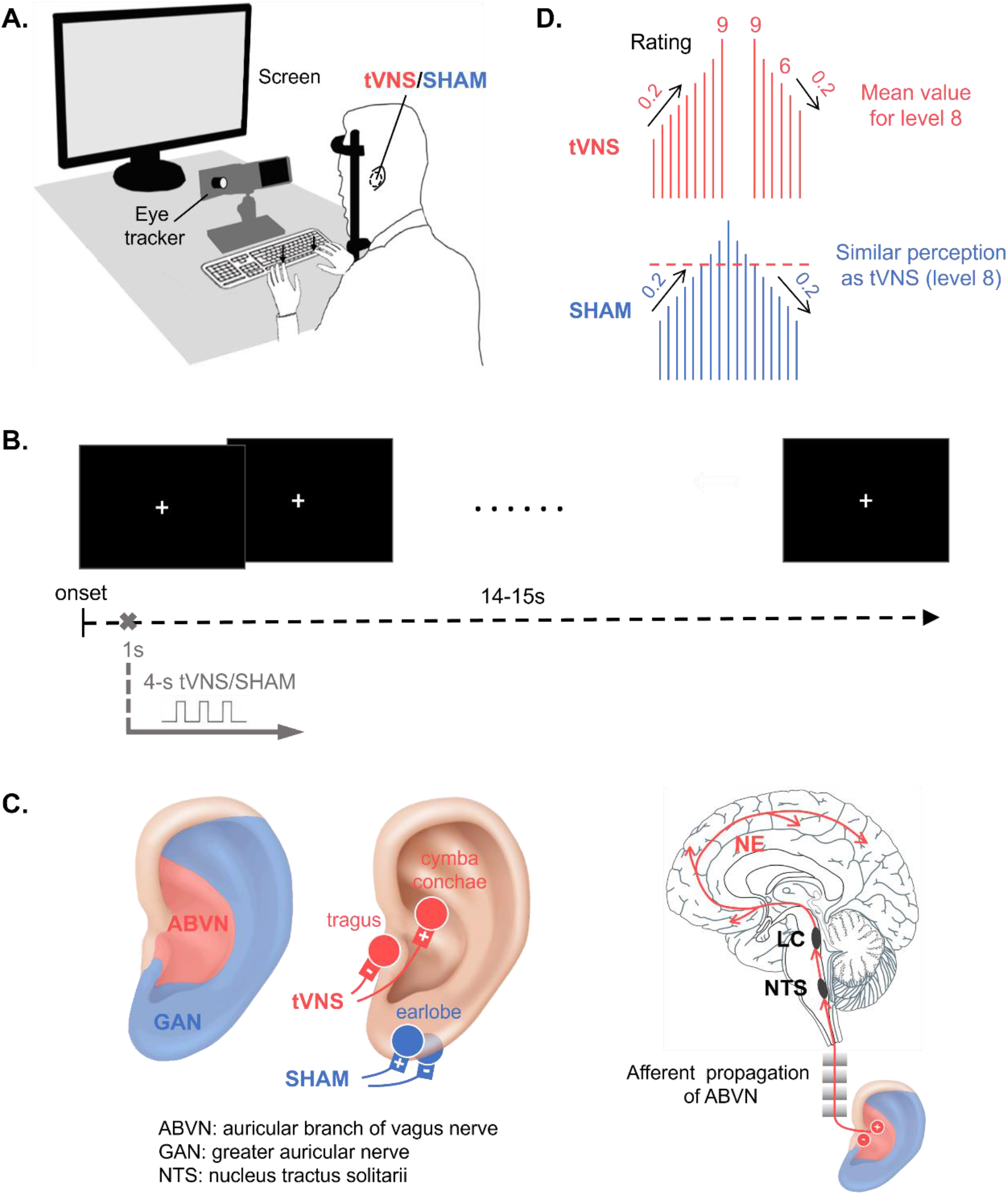
Experimental setup and design in Experiment 1. **A. Experimental setup**. Subjects sat in front of a computer screen, with their head positioned on a chin-rest apparatus (60cm from the monitor). The screen (operating at 60Hz and a resolution of 1360 × 768 pixels) displayed a central cross (Experiment 1) or the RDM task (Experiment 2). Subjects rested their left and right forearms on the table (Experiment 1) or placed the left and right index fingers on the F12 and F5 keys of an upside-down keyboard, respectively (Experiment 2). Pupillometry was recorded throughout the session in both experiments. Subjects were instructed to minimize movements and eye blinks during the recording. **B. Trial illustration in Experiment 1**. Participants were instructed to focus on a white cross displayed in the center of a black screen while receiving tVNS/SHAM stimulation every 14-15s at rest. **C. Electrode montage in tVNS and SHAM conditions**. Electrodes (Axiothera, German) were put on the cymba conchae and tragus of the left ear in tVNS condition (secured with medical paste for optimal contact). This allows to generate an electric field covering the area innervated by the auricular branch of the vagus nerve, thereby specifically activating the vagal afferent pathway up to the nucleus tractus solitarii (NTS), which in turn transmits impulses to locus coeruleus (LC) and activates norepinephrine (NE) neurons that project throughout the brain. Electrodes were put on the left earlobe in SHAM condition. This allows to generate an electric field covering the area of the greater auricular nerve whose stimulation is not expected to induce brainstem or cortical activation. **D. Stimulation intensity calibration**. Subjects underwent a short “Method of Limits” procedure to select the maximal comfortable tVNS intensity. Subjects received increasing and decreasing series of 4s tVNS trials and rated their subjective sensation for each stimulation on a 10-point scale, ranging from no sensation [0], light tingling [3], strong tingling [6], to painful [10]. The intensity always started at 0.1mA and increased by 0.2mA until subjects reported a sensation of 9. Before starting the decreasing series, the same intensity was repeated. Then, it was reduced trial by trial in 0.2mA steps until a subjective sensation of 6 or below was experienced. The final intensity for tVNS was calculated based on the average of the intensities rated as 8. To match the perception in the two conditions, intensity in the SHAM condition was set to the equal perception of the final intensity in tVNS condition (rated as 8). Specifically, the experimenter modulated the intensity of SHAM with increasing and decreasing series in 0.2mA steps, and subjects were asked to report how they felt after each SHAM stimulation without rating. The final intensity for SHAM was calculated based on the average of intensities felt as similar to the final intensity for tVNS rated as 8. In Experiment 1, average intensities were 1.63 ± 1.23mA for tVNS and 5.85 ± 3.12mA for SHAM; in Experiment 2, the values were 1.42 ± 0.61mA and 3.63 ± 1.81mA, respectively.

### Experiment 1: Validating the transcutaneous vagus nerve stimulation (tVNS) protocol

#### Procedures at rest

Previous studies on tVNS have used various protocols to activate LC, resulting in mixed effects on pupil size [57, 58]. Most of these studies applied tVNS for extended periods to induce long-term effects [59-62], but fewer have explored brief tVNS bursts lasting a few seconds for transient effects [63-65]. Experiment 1 aimed to validate a protocol using a brief 4s tVNS train in participants at rest. Participants were instructed to focus on a white cross on a black screen (see Fig. 1B). Pulses were generated using a Digitimer (Digitimer Ltd., model DS3), triggered by a Master8 device (A.M.P.I., model Master-8). Surface electrodes delivered pulses to the skin of the left ear (see Fig. 1C). In tVNS condition, an anode electrode was placed on the left cymba conchae, while the cathode was taped to the left tragus [48, 49]. Such a montage allows to stimulate the auricular branch of the vagus nerve [42, 50, 51], thereby activating the LC [41, 52, 53]. In SHAM condition, electrodes were attached to the left earlobe, which is not expected to activate the brainstem or cortex [54-56]. Pulses (200 µs) were delivered at 25Hz for 4s in both conditions. The experiment always started with a calibration session during which the tVNS intensity was set just below the painful threshold, while SHAM was adjusted to match the perceived intensity of tVNS (see Fig. 1D for details on intensity calibration). During the main part of Experiment 1, each participant completed 4 to 6 blocks of 20 trials with either tVNS or SHAM stimulation every 14-15s at rest. More details on the procedures in this experiment are provided in the Supplementary Materials.

#### Pupillometric data acquisition and analyses

Pupil size was recorded at 1000 Hz using an Eyelink 1000+ eye-tracker (SR Research) and analyzed in Matlab. After data cleaning and artifact rejection (see Supplementary Materials), we excluded 2 subjects. The remaining subjects (n=20) had an average of 91 ± 13.7 trials for pupil analyses. Pupil data were segmented into 9s-trials spanning from 1s before to 8s after the stimulation onset. To avoid arbitrary units, we converted pupil size into z scores computed across all conditions in each subject. Pupil baseline was defined as the average size from -1 to 0s relative to stimulation onset, and pupil dilation was calculated as the change from baseline at each time point [35, 66]. Pupil dilation under tVNS and SHAM were compared using a cluster-based paired t-test with Monte-Carlo permutations [67, 68]. To quantify differences, we identified a window of full duration at half-maximum (FDHM) from the grand average of tVNS and SHAM [54]. Pupil dilation within this FDHM window was averaged and analyzed using a linear mixed-effects model (Jamovi) with Block-Type (tVNS, SHAM) as the factor. The same model was used to compare pupil baseline between tVNS and SHAM blocks.

### Experiment 2: Applying tVNS during perceptual decision-making

#### Procedures using the random dot motion (RDM) task

Here, we aimed to test the causal role of LC in perceptual decision-making using the same tVNS procedure established in Experiment 1. We used a variation of the RDM task where participants had to press a key (on an inverted keyboard; see Fig. 1A) with the left or right index finger as quickly and accurately as possible depending on whether coherently moving dots progressed to the left or right, respectively. As shown in Figure 2A, each experiment started with a training session. Then, we performed two calibration sessions: a first session served to set the individual coherence (c’) value, adjusted to obtain an accuracy of approximately 70% (c’ value = 0.2 ± 0.15; see Fig. 2B for individual values), while the second one was to determine the personalized tVNS/SHAM intensity (see details in Fig. 1D). Then during the main experiment, subjects performed the RDM task across 8 blocks of 40 trials, split into 4 tVNS and 4 SHAM blocks, with rest breaks in between. These blocks involved a 4-second tVNS/SHAM train on each trial applied 3.5s after trial onset and ending either before or at the onset of coherent motion (see Fig. 2C). More details on procedures in this experiment are provided in Figures 1 and 2 as well as in the Supplementary Materials.

**Figure 2.**
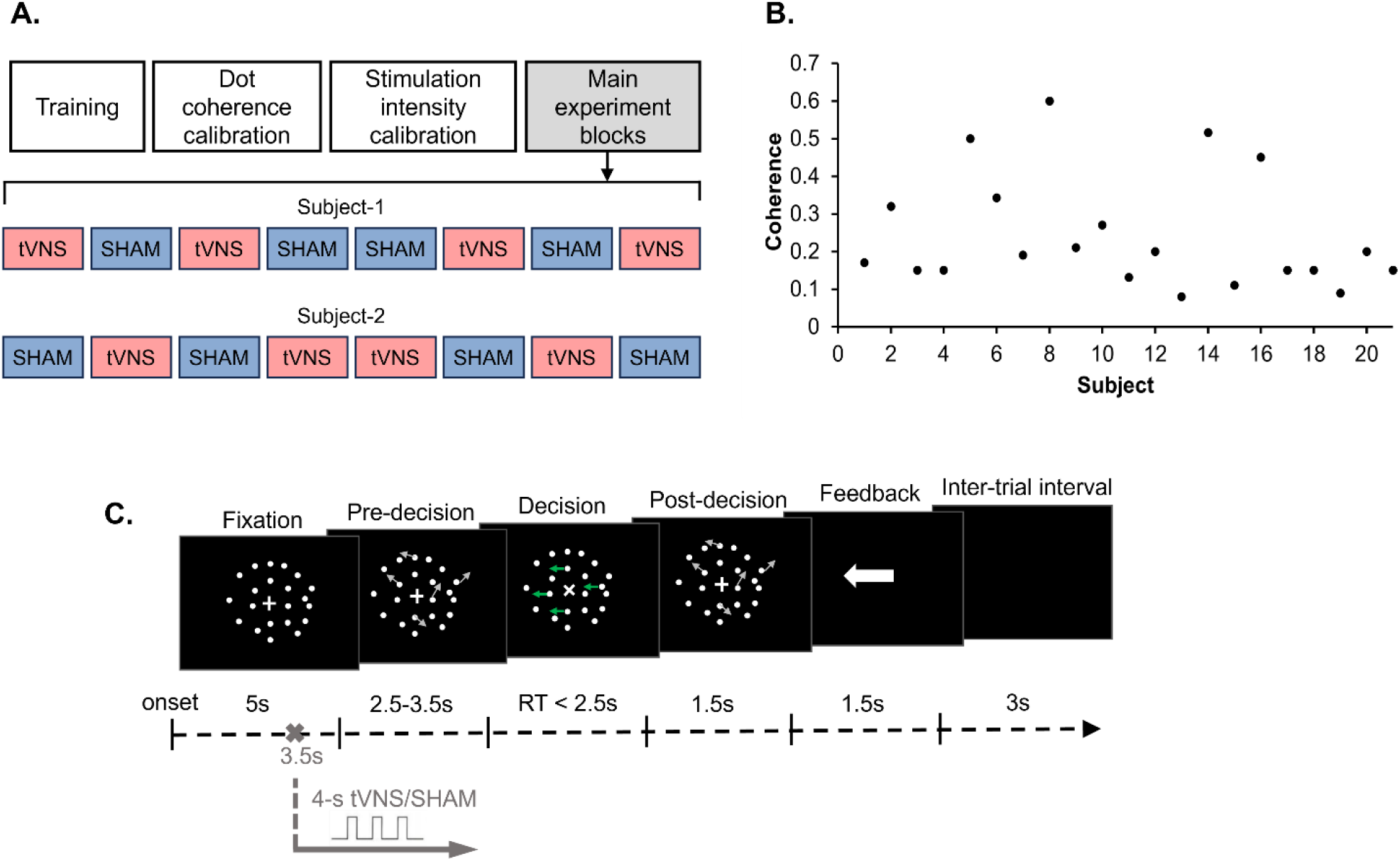
Experimental design in Experiment 2. **A. Session design in Experiment 2**. Each subject underwent a training and two calibration sessions (dot coherence and stimulation intensity) before they got involved in the 8 main experiment blocks. The tVNS and SHAM blocks were alternated, and the sequence of the first 4 blocks was reversed for the second set. Block sequences were counterbalanced across subjects, as shown in two representative subjects. **B. Individual coherence c’ in Experiment 2**. The proportion of dots moving in a coherent direction (c’) was determined at the beginning of the experiment for each subject to ensure an accuracy of 70%. It ranged from 0.08 to 0.6. **C. Trial illustration in Experiment 2**. In the random dot motion (RDM) task, each trial started with a Fixation phase (5s) during which subjects were instructed to maintain fixation on a centrally presented cross surrounded by stationary dots. In the subsequent Pre-decision phase, dots started to move randomly at a speed of 5 degrees/s. The duration of this phase was drawn from a uniform distribution with a range of 2.5-3.5s. The transition from stationary to randomly moving dots was designed to avoid non-task-related changes in pupil size caused by sudden luminance increments. Then came the Decision phase, during which the central cross “+” changed to an “x” and a proportion of dots started to move in a coherent direction (leftward or rightward). In concrete terms, dots in the first three frames were repositioned after two subsequent frames. The new location of a dot was either random or in the dominant direction of motion on that trial (as represented by green arrows: leftward motion in this example). The proportion of coherently moving dots in each trial was defined as coherence c’. Coherent dot motion directions were equiprobable and randomly selected across trials. Subjects were required to indicate the direction of the coherent dot motion by pressing on a key with their left or right index finger (F12 or F5 key on an inverted keyboard, respectively). Here, we set a time limit for the decision at 2.5s to make sure subjects were fully engaged in the task. Upon response execution, coherent dot motion transitioned to purely random dot motion (Post-decision phase; 1.5s) to minimize post-decisional sensory evidence accumulation [69]. Subjects then received feedback (1.5s) in the form of a white arrow pointing to the left or right to show the actual motion direction in that trial. If subjects had not responded within the specified time (2.5s), the arrow was replaced by the caption ‘missed!’. A final intertrial phase of 3s followed, during which a black background was displayed. In each trial, a 4-second tVNS/SHAM train was applied 3.5s after trial onset and ending either before or at the onset of coherent motion.

#### Behavioral measures and analyses

We used Reaction Time (RT) and Accuracy to measure performance in the RDM task. RT was the time from coherent motion onset to key press, and Accuracy was the percentage of correct responses. Trials with no response (< 2.3%) or very fast responses (< 500ms, < 0.3%) were excluded. Consequently, behavioral analyses were performed on an average of 273 ± 58.7 trials in Experiment 2 (n = 21). Trials were categorized by Block-Type (tVNS, SHAM), and some analyses also separated trials based on whether they involved a correct choice or an error (Trial-Success: correct, error) and/or whether they followed a correct choice or an error in the preceding trial (Trial-History: after-correct, after-error). Depending on the dot motion coherence (c’ value used in the RDM task), subjects were given a Coherence-Rank (1 to 16, subjects with the same c’ values were attributed as the same rank) and split into two Coherence-Groups: low-coherence (n = 11) and high-coherence (n = 10). Statistical analyses were conducted with Jamovi using linear (RT) or generalized (Accuracy) mixed-effect models. The main factor was Block-Type, with additional factors including Trial-Success and Trial-History, as well as individual Coherence-Rank as a covariate. Post-hoc tests with Holm-Bonferroni corrections were conducted to address specific interactions, and JASP was used for Pearson’s Partial correlations. Results were expressed as means ± SEM, with an alpha level of 0.05.

#### Pupillometric data acquisition and analyses

Pupil size was recorded and preprocessed using the same method as in Experiment 1, leading to the exclusion of 1 subject (n=20). Additionally, after excluding trials with no response or very fast responses, an individual average of 271 ± 59.6 trials were included for pupil analyses. Here too, Pupil data were segmented into 9-second trials, converted into z-scores across all conditions, with pupil baseline assessed in the -1 to 0s relative to stimulation onset and pupil dilation calculated as the change from baseline at each time point [35, 66]. To assess how tVNS influenced LC-NE activity during the RDM task, we analyzed pupil dilation aligned to either stimulation onset ([-1 8]s) or coherent motion onset ([-1 4]s). We then compared pupil dilation using paired t-tests (Block-Type: tVNS, SHAM) or repeated-measures ANOVAs (considering also Trial-Success: correct, error; Trial-History: after-correct, after-error, and Coherence-Group: low-coherence, high-coherence) with cluster-based Monte-Carlo permutations.

#### Drift diffusion model

We considered the DDM to explore the computational mechanisms underlying the behavioral effects. The DDM provides three key parameters: drift rate (reflecting the efficiency of sensory evidence accumulation) [45], boundary (representing decision-making criteria) [70], and non-decision time (capturing processes other than decision-making, like motor execution) [71]. Here, we used a refined DDM with linearly collapsing boundaries, where a lower boundary intercept indicates a higher initial urgency, and a steeper boundary slope signifies increasing urgency over time. These parameters were estimated by fitting RT and Accuracy data from the RDM task for 16 participants based on Block-Type (tVNS, SHAM) and Trial-History (after-correct, after-error). Details on parameter fitting and recovery analysis are in the Supplementary Materials. Linear mixed-effects models were used to compare parameters across Block-Type, Trial-History, and Coherence-Rank.

## Results

### Experiment 1

#### tVNS increased pupil dilation at rest

In Experiment 1, pupil size was measured at rest in both tVNS and SHAM blocks (n=20). As shown in Figure 3A, tVNS elicited a rapid pupil dilation, peaking (0.22) at 1.50s, and returning to baseline by 4.23s. SHAM stimulation also induced pupil dilation but with a smaller peak (0.11) and quicker return to baseline (3.46s). A paired t-test with cluster-based permutations showed significantly greater pupil dilation under tVNS from 0.78s to 3.12s, and from 3.97s to 5.22s post-stimulation (p < 0.05). A linear mixed-effects model on the full duration at half-maximum (FDHM; 0.45-2.44s) showed significantly higher pupil dilation in tVNS (0.18 ± 0.03) compared to SHAM (0.08 ± 0.02) blocks (F(1, 19) = 16.30, p < 0.001, Fig. 3B). There was no significant difference in pupil baseline between tVNS and SHAM blocks (F(1, 38) = 1.73, p = 0.20, Fig. 3C).

**Figure 3.**
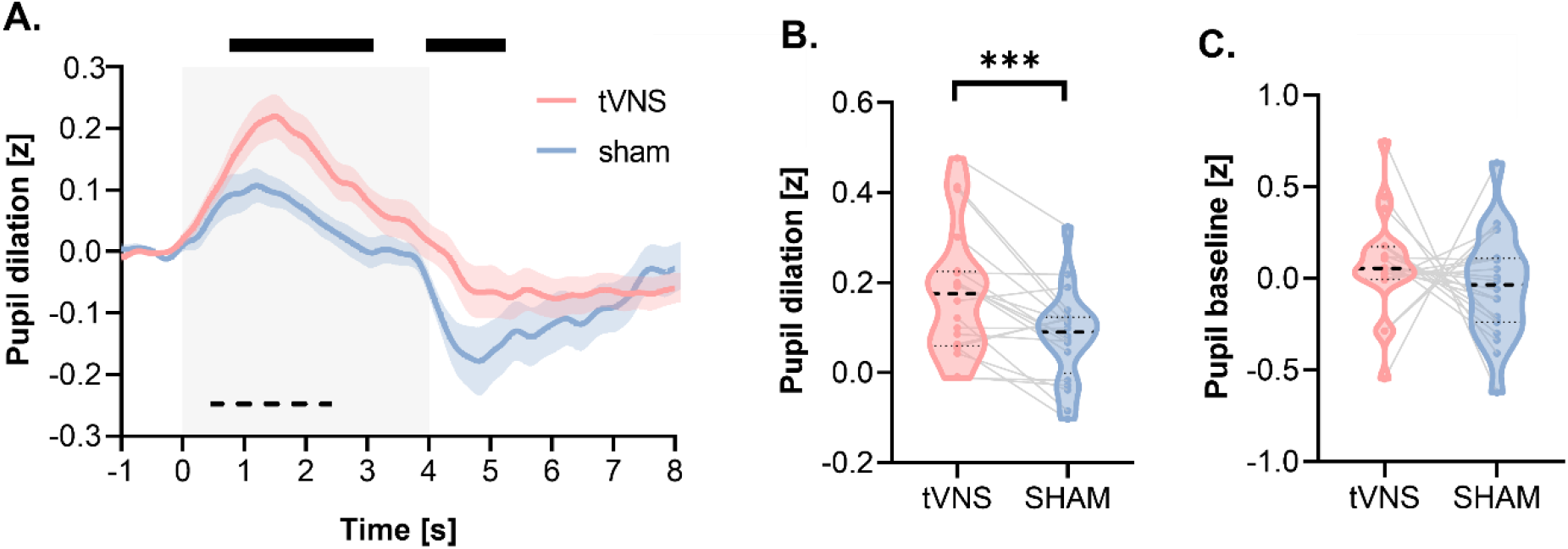
Effect of tVNS on pupil data at rest (Experiment 1, n=20). The graphs display pupil data, separately for tVNS (red) and SHAM (blue) blocks. **A. Grand average of pupil dilation**. Pupil dilation is aligned to stimulation onset [-1 8]s, with shaded areas around the two traces indicating the SEM in tVNS and SHAM blocks. The gray transparent rectangle indicates the 4 s-stimulation period. The top black lines indicate statistical significance in pupil dilation between tVNS and SHAM blocks using paired t-test with cluster-based permutations (p < 0.05 from 0.78s to 3.12s, and from 3.97s to 5.22s post-stimulation onset). The dashed black line indicates the full duration at half-maximum (FDHM, 0.45-2.44s), which is used to compute the average pupil dilation in each subject. **B. Pupil dilation during the FDHM**. Note the greater pupil dilation in tVNS compared to SHAM blocks, as evident from the shift in median values (black dotted lines), distributions as violin plots, and the consistent effect on individual pupil data (thin gray lines). **C. Pupil baseline**. Note the comparable pupil baseline in tVNS and SHAM blocks. ***: *p* < 0.001.

### Experiment 2

#### tVNS enhanced accuracy with no effect on RT in the RDM task

Experiment 2 investigated the effect of tVNS on decision-making performance in the RDM task (n=21). Interestingly, tVNS significantly enhanced accuracy, with participants making more correct decisions under tVNS (78.2 ± 0.78%) compared to SHAM (75.6 ± 0.80%), as shown by a significant Block-Type effect in the generalized mixed-effects model (*χ*^2^ (1) = 5.06, p = 0.024; see Fig. 4A). However, tVNS did not affect decision speed: RTs were comparable in tVNS (1.3 ± 0.08s) and SHAM (1.3 ± 0.07s) blocks, with no significant Block-Type effect in the linear mixed-effects model (F(1, 18.9) = 0.25, p = 0.620; see Fig. 4B). Therefore, it appears that enhancing LC-NE activity through tVNS improved decision accuracy without changing decision speed.

**Figure 4.**
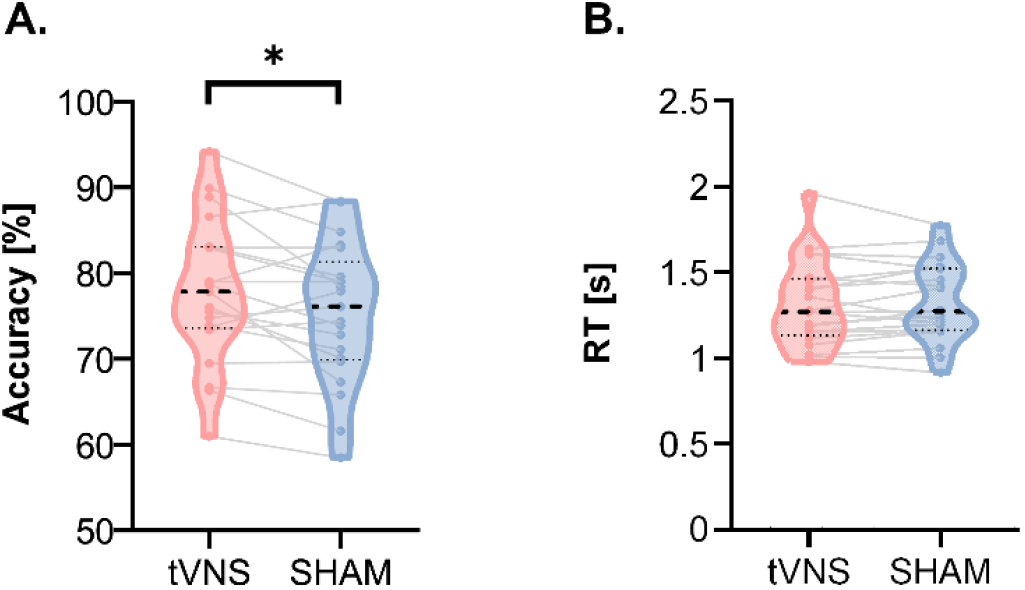
Effect of tVNS on behavior in the RDM task (Experiment 2, n=21). The graphs display behavioral data, separated for tVNS (red) and SHAM (blue) blocks, with median values featured as black dotted lines, distributions as violin plots, and individual data as thin gray lines. **A. Accuracy**. Note the greater accuracy in tVNS compared to SHAM blocks. **B. Reaction time (RT)**. Note the comparable RTs in the two block types. *: *p* < 0.05.

#### tVNS increased pupil dilation during the RDM task

Experiment 2 also assessed the effect of tVNS on pupil dilation during the RDM task (n=20). When aligned to stimulation onset (Fig. 5A), pupil size increased during the task, especially at the onset of random dot motion in the Pre-decision phase. It peaked at 5.13s in tVNS blocks (peak dilation: 0.66) and at 5.10s in SHAM blocks (peak dilation: 0.59), around the time participants made their responses. Pupil size then returned to baseline around 7.85s. Paired t-tests with cluster-based permutations showed significantly greater pupil dilation under tVNS from 1.2s to 5.8s post-stimulation (p < 0.05). When aligned to coherent motion onset, pupil dilation was also larger in tVNS blocks for up to 1.2s after onset (p < 0.05; Fig. 5B). This time nearly coincided with RTs (about 1.3s), indicating that tVNS effect on pupil size covered most of the decision phase, even though the stimulation ended earlier.

**Figure 5.**
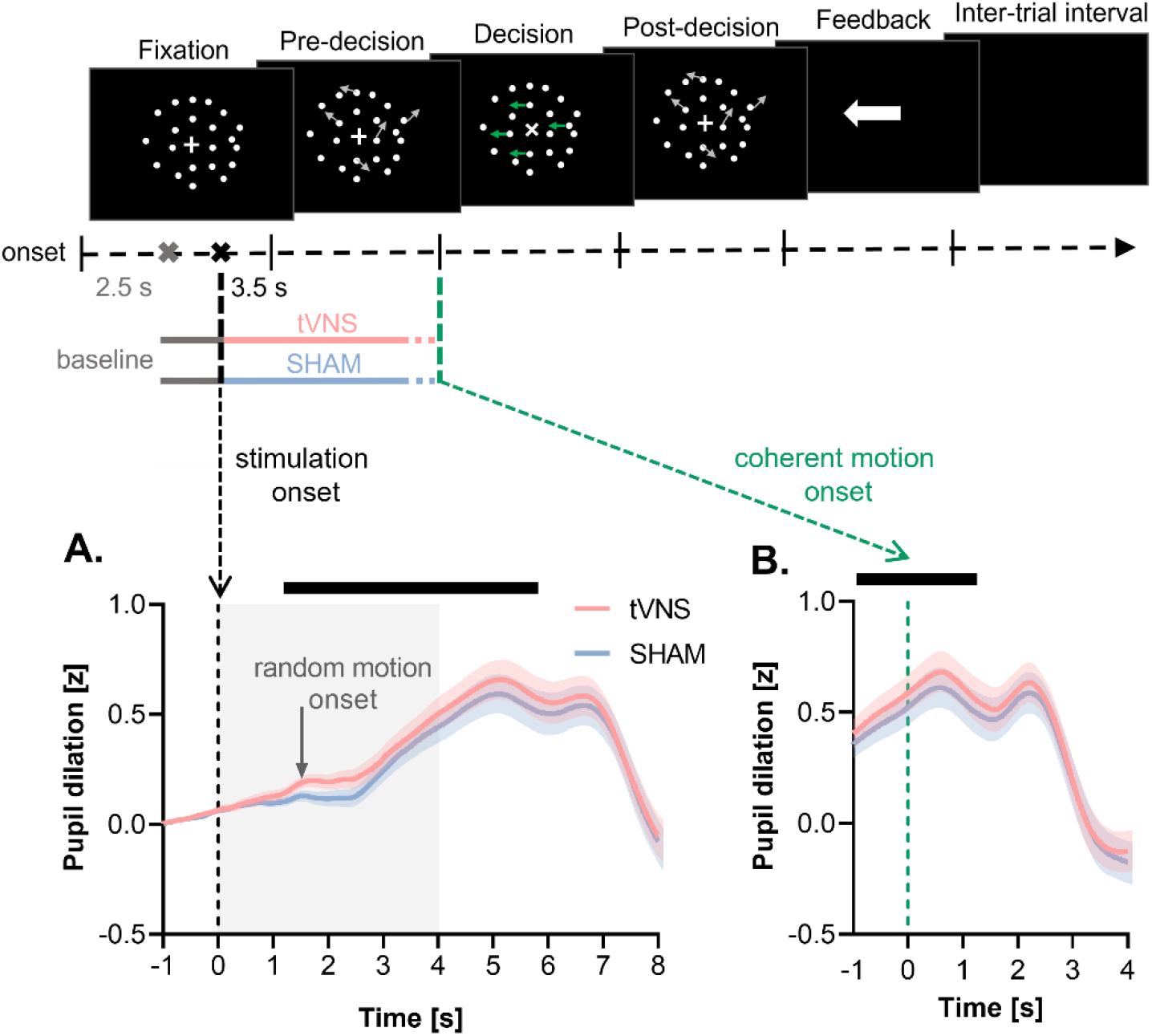
Effect of tVNS on pupil dilation in the RDM task (Experiment 2, n=20). The graphs display pupil dilation separately for tVNS (red) and SHAM (blue) blocks, with shaded areas around the traces indicating the SEM in both block types. The top black lines in both graphs indicate statistical significance using paired t-test with cluster-based permutations. **A. Pupil dilation aligned to stimulation onset**. Note the increase in pupil dilation that becomes significantly greater in tVNS than SHAM blocks from 1.2s to 5.8s post-stimulation onset. **B. Pupil data aligned to coherence motion onset**. Note the greater pupil dilation in tVNS than SHAM blocks up to 1.2s post-coherent motion onset.

#### tVNS did not alter pupil baseline, but the latter showed a strong increase after errors

To examine whether tVNS-induced pupil changes extended into the intertrial interval, we analyzed pupil baseline with a linear mixed-effects model. Here, we included the factor of Block-Type, as well as Trial-History, as errors are known to increase pupil size in subsequent trials [72]. There was no significant effect or interaction involving Block-Type (F < 1.59, p > 0.206, Fig. 6A), suggesting that the 10-13s inter-stimulation interval was long enough to reset baseline LC-NE activity. However, pupil baseline was significantly influenced by Trial-History, with higher baselines following errors compared to correct trials (F(1, 5401.1) = 72.50, p < 0.001, Fig. 6B), indicating increased LC-NE activity after errors.

**Figure 6.**
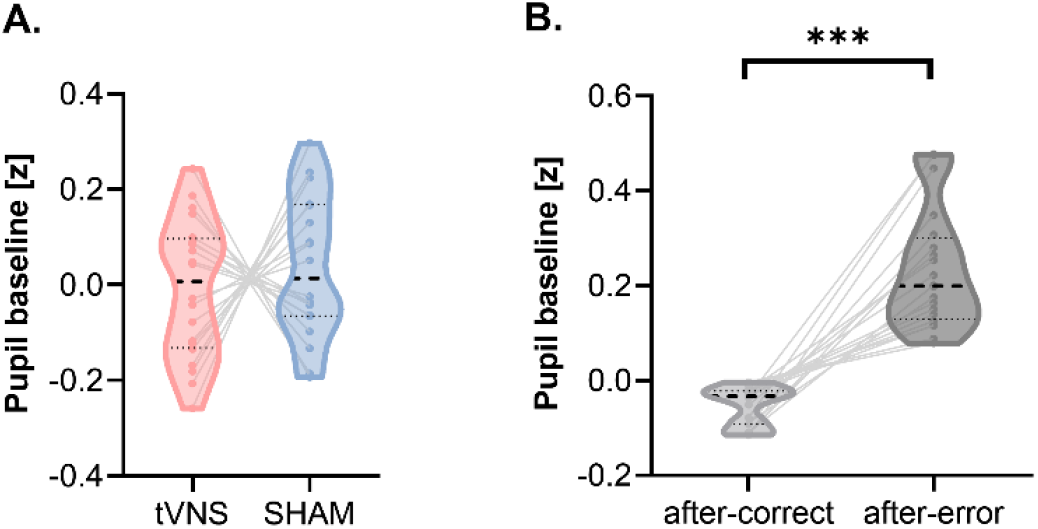
Pupil baseline in the RDM task (Experiment 2, n = 20). The graphs display median values as black dotted lines, distributions as violin plots, and individual data as thin gray lines. **A. Block-Type effect**. Note the comparable pupil baseline in tVNS (red) and SHAM (blue) blocks. **B. Trial-History effect**. Note the greater pupil baseline in after-error (dark gray) than after-correct trials (light gray). ***: *p* < 0.001.

#### tVNS enhanced accuracy specifically after errors

We conducted a follow-up analysis to see if the effect of tVNS on behavior depended on Trial-History, given its impact on baseline LC-NE activity. Here, we also considered Coherence-Rank, since coherence levels may affect participants’ responses to errors [73, 74] and indicate task proficiency: participants with lower coherence levels needed less sensory evidence to achieve 70% accuracy, indicating higher task proficiency.

For Accuracy, the generalized mixed-effects model showed significant main effects of Block-Type (*χ*^2^ (1) = 6.29, p = 0.012) and Trial history (*χ*^2^ (1) = 6.30, p = 0.012), along with a three-way interaction involving Coherence-Rank (*χ*^2^ (2) = 9.84, p = 0.007). To explore this interaction, we considered participants with Coherence-Groups (low-coherence, high-coherence). As shown in Figure 7A, tVNS had no effect on more proficient subjects (low-coherence group): Accuracy was similar between tVNS and SHAM blocks, for both after-errors (Z = 0.39, p_holm-bonferroni_ = 1) and after-correct responses (Z = 2.32, p_holm-bonferroni_ = 0.081). In contrast, for less proficient subjects (high-coherence group), tVNS significantly improved accuracy after errors (Z = 3.139, p_holm-bonferroni_ = 0.006) but not after correct responses (Z = -0.94, p_holm-bonferroni_ = 0.698). Notably, this group showed a marked accuracy decline following errors in SHAM blocks (Z = 5.06, p_holm-bonferroni_ < 0.001), which was not observed in the low-coherence group (Z = -0.38, p_holm-bonferroni_ = 1). This suggests that tVNS mitigated the accuracy decline following errors in the high-coherence group. Specifically, participants who showed a larger accuracy decline after errors in SHAM blocks benefited more from tVNS in terms of accuracy improvement (R = -0.536, p = 0.015, see details in Supplementary Fig. 1A).

**Figure 7.**
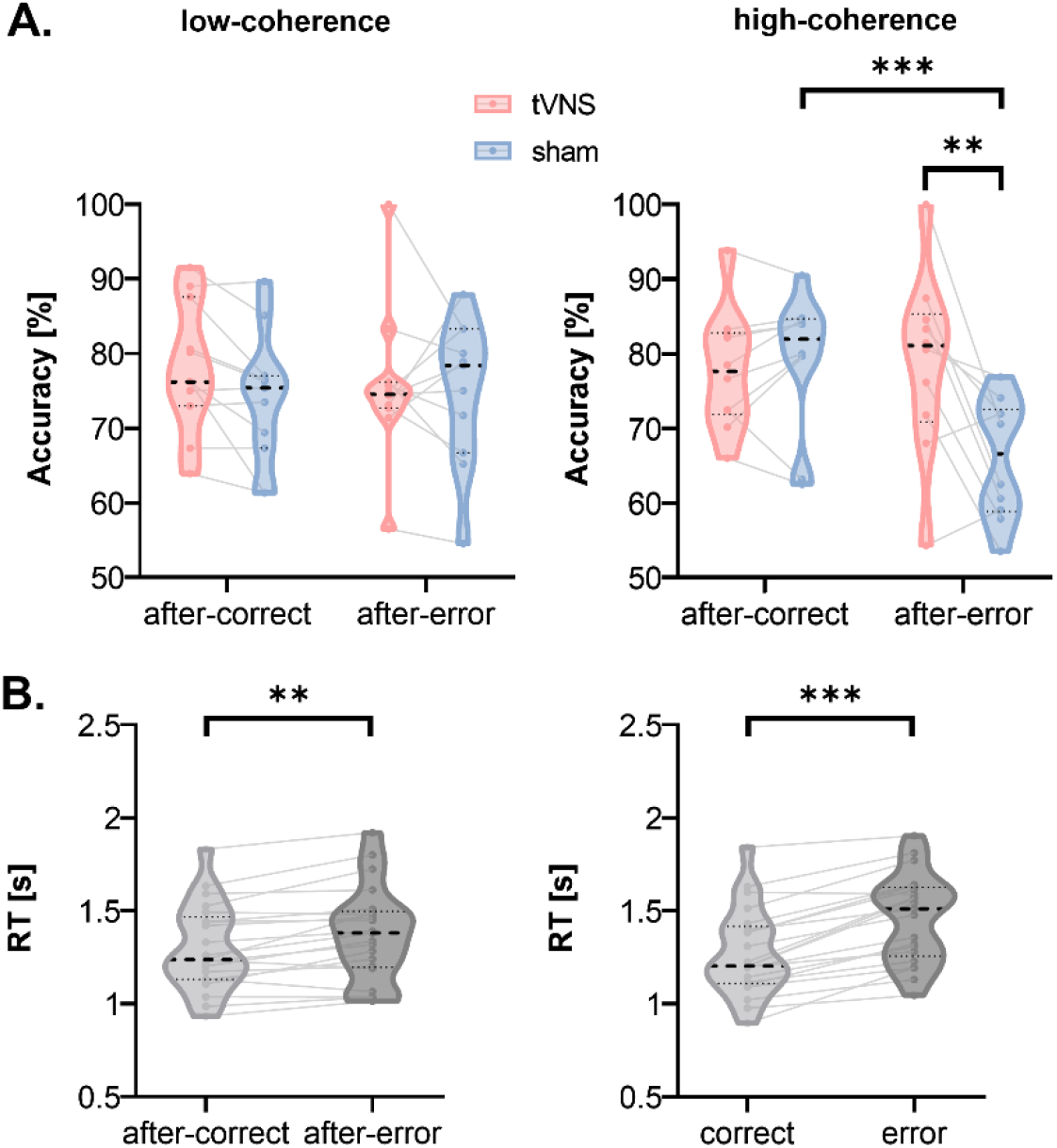
Effect of tVNS on behavior in the RDM task, as a function of Trial-History and Coherence-Rank (Experiment 2, n=21). **A. Accuracy**. The accuracy data features median values as black dotted lines, distributions as violin plots, and individual data as thin gray lines, separated for tVNS (red) and SHAM (blue) blocks. tVNS effect on accuracy, showing no impact on the low-coherence group regardless of Trial-History (left panel), whereas significantly improved accuracy in the high-coherence group, specifically in trials following errors (right panel). **B. Reaction Time (RT)**. The RT data features median values as black dotted lines, distributions as violin plots, and individual data as thin gray lines. tVNS effect on RT, showing slower RTs in after-error (dark gray) compared to after-correct (light gray) trials (left panel), and longer RTs when made errors (dark gray) than correct (light gray) responses (right panel). **: *p* < 0.01. ***: *p* < 0.001.

For RTs, the linear mixed-effects model, including Block-Type (tVNS, SHAM), Trial-History (after-correct, after-error), and Trial-Success (correct, error) as factors, as well as Coherence-Rank as a covariate, revealed no significant main effect or interaction involving Block-Type (F < 2.11, p > 0.105). If anything, participants exhibited slower RTs following errors compared to after-correct responses (F(1, 23.1) = 12.39, p = 0.002, Fig. 7B, left panel), consistent with post-error slowing literature [75-77]. Additionally, errors were associated with longer RTs than correct responses (F (1, 22.5) = 555.45), p < 0.001, Fig. 7B, right panel), reflecting the common pattern where slower decisions are linked to less accumulated evidence, increasing the likelihood of incorrect responses [78, 79]. As expected, there was no correlation between post-error slowing in SHAM blocks and any potential tVNS effect on RTs after errors (R = -0.262, p = 0.264, see details in Supplementary Fig. 1B)

#### tVNS effect on pupil dilation during the RDM task did not depend on trial history or coherence

Given the selective behavioral effect of tVNS after errors, especially in the high-coherence group, we conducted an additional analysis to see if pupil dilation was more pronounced after errors in this group. We performed a repeated-measures ANOVA with cluster-based permutations on pupil dilation aligned to stimulation, considering Block-Type, Trial-History, and Coherence-Group as factors. However, none of the interactions involving Block-Type were significant (p > 0.05), indicating that tVNS increased pupil dilation uniformly across conditions. Therefore, the selective effect of tVNS on behavior is not reflected in the magnitude of pupil dilation. In other words, while pupil dilation may index LC-NE activation, it does not seem to directly reflect the extent to which tVNS enhances behavior in this study.

#### tVNS increased the drift rate specifically after errors

We investigated the effect of tVNS on DDM parameters: drift rate, boundary intercept, boundary slope, and non-decision time. For drift rate, the linear mixed-effects model revealed a triple interaction between Block-Type, Trial-History, and Coherence-Rank (F(2, 42) = 3.60, p = 0.036). In the high-coherence group, the drift rate was significantly lower for after-error compared to after-correct trials in SHAM blocks (t(21) = 3.94, p_holm-bonferroni_ < 0.001, Fig. 8A, right panel). However, tVNS increased the drift rate after errors compared to SHAM (t(21) = 2.65, p_holm-bonferroni_ = 0.045), eliminating the Trial-History effect (t(21) = - 0.16, p_holm-bonferroni_ = 0.871). Interestingly, no effects were found in the low-coherence group (F < 0.12, p > 0.26, Fig. 8A, left panel), aligning with the accuracy results.

**Figure 8.**
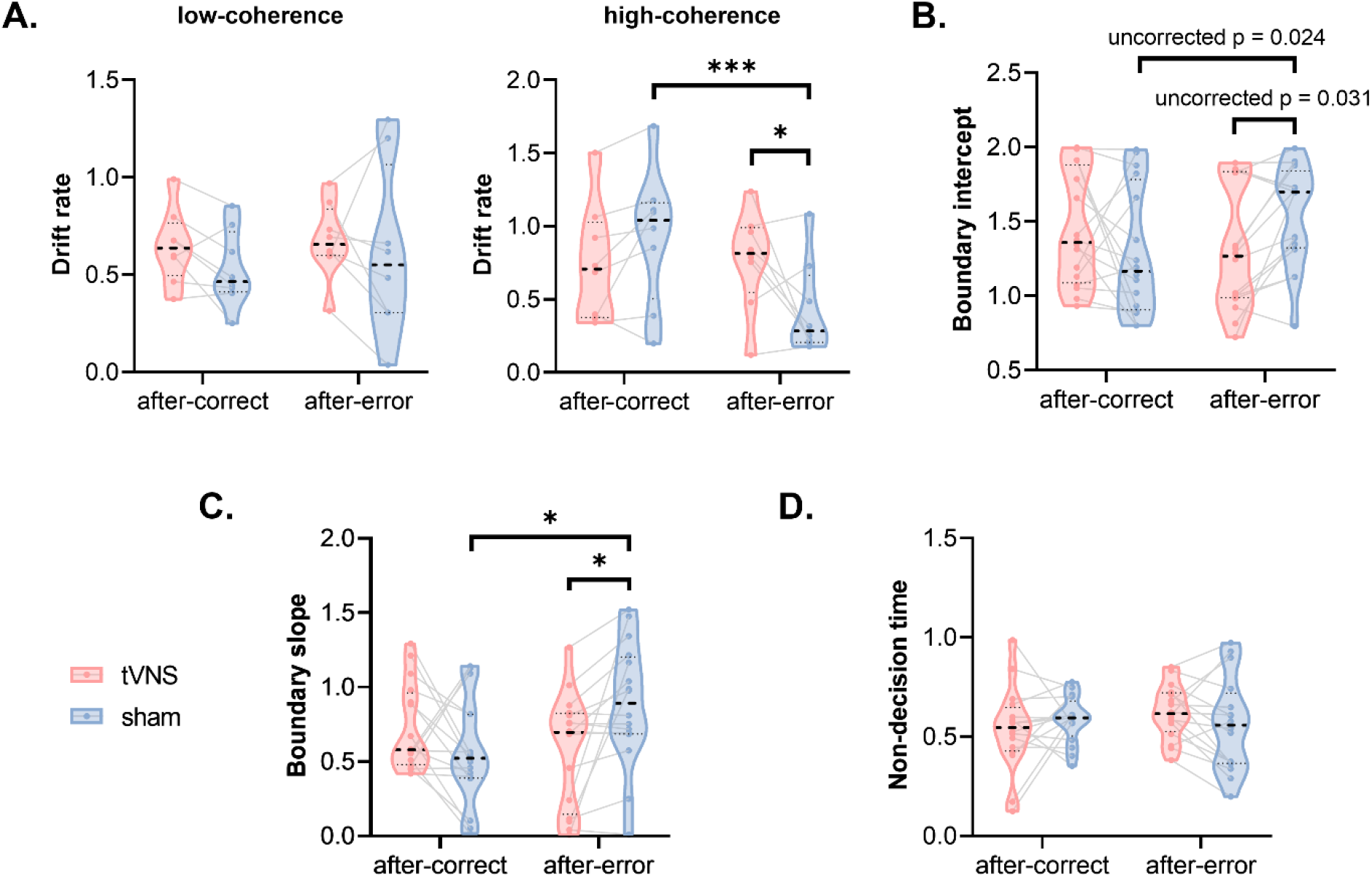
tVNS effect on Drift Diffusion Model (DDM) parameters (Experiment 2, n=16). The graphs display DDM data, separated for tVNS (red) and SHAM (blue) blocks, with median values featured as black dotted lines, distributions as violin plots, and individual data as thin gray lines. **A. Drift rate**. For the high-coherence group selectively (right panel), note the tVNS-induced increase in drift rate for after-error trials that occurs in parallel to the after-error drop in drift rate evidenced in SHAM blocks. **B. Boundary intercept**. Note the marginal tVNS-induced decrease in boundary intercept for after-error trials that occur in parallel to the marginal after-error increase in boundary intercept in SHAM blocks. **C. Boundary slope**. Note the lower boundary slope in after-error trials under tVNS compared to SHAM blocks where the boundary slope increases in after-error compared to after-correct trials. **D. Non-decision time**. Note the consistent values across conditions, reflecting no tVNS effect on non-decision times. *: *p* < 0.05. ***: *p* < 0.001.

For boundary intercept and slope, we both found significant interactions between Block-Type and Trial-History (intercept: F(1, 42) = 5.01, p =0.031; slope: F(1, 42) = 5.29, p = 0.031). As shown in Figures 8B and 8C, participants tended to exhibit a higher intercept and displayed a steeper slope after errors compared to correct responses in SHAM blocks (intercept: t(42) = -2.35, p = 0.024; slope: t(42) = -3.1, p_holm-bonferroni_ = 0.012). However, tVNS tended to lower the intercept and decreased the slope after errors compared to SHAM blocks (intercept: t(42) = -0.23, p = 0.031; slope: t(42) = -2.91, p_holm-bonferroni_ = 0.018), eliminating the Trial-History effect in tVNS blocks (intercept: t(42) = 1.25, p_holm-bonferroni_ = 0.217; slope: t(42) = 1.36, p_holm-bonferroni_ = 0.183). Finally, there were no significant effects for non-decision time (F < 3.06, p > 0.080, see Fig. 8D).

We also calculated Delta values (after-error minus after-correct) for these parameters to assess specifically how post-error adjustments are affected by tVNS. These data are described in the Supplementary Materials, both in the text and in Supplementary Figure 2.

## Discussion

In this study, we applied tVNS during the RDM task to explore the causal role of LC in decision-making. We found that tVNS improved decision accuracy (but not RT) and increased drift rate, particularly following errors and in participants prone to accuracy drops after errors. Pupil data confirmed effective LC stimulation by tVNS. These results support the “gain” hypothesis, suggesting that LC-NE activity facilitates decision processes by enhancing the signal-to-noise ratio in perceptual processing [24].

A key observation in this study is that tVNS improved accuracy in the RDM task without affecting RTs, enabling participants to make more accurate decisions at the same speed. These behavioral findings, derived from a causal approach, align with the well-established “gain” hypothesis from animal research.

However, this raises the question of how to reconcile such a view with earlier correlational studies in humans, which proposed an “urgency” function for the LC, based on the observation of greater pupil dilation during fast (less accurate) decisions compared to slower (more accurate) ones [35]. In those studies, fast decisions were typically driven by strict time constraints, leading to the plausible hypothesis that elevated LC-NE activity was observed there because it played a direct role in generating urgency for the acceleration of behavior to meet deadlines, albeit at the cost of accuracy [38, 80]. However, more consistent with our findings, it is also plausible that enhanced LC-NE activity in such scenarios helps manage the challenge of making fast decisions without sacrificing too much accuracy. In this view, LC recruitment under an urgent deadline might not generate urgency per se but rather help to maintain accuracy under time pressure, as previously suggested by the positive relationship between pupil dilation and accuracy in such situations [37, 81], including a recent study from our lab [39]. Therefore, our current tVNS findings prompt us to reinterpret previous human studies through the lens of “gain” hypothesis [9, 13, 82].

Here, the beneficial effect of tVNS on accuracy was most pronounced in trials following errors, especially for less proficient participants who needed higher coherence levels in dot motion to reach 70% target accuracy. Interestingly, these participants exhibited a marked decrease in accuracy after errors during SHAM blocks, a decline that disappeared in tVNS blocks. In contrast, more proficient participants, who operated at lower coherence levels, did not experience a post-error drop in accuracy, and tVNS did not impact their accuracy. This suggests that tVNS selectively influenced behavior in a particular subset of trials and participants associated with compromised performance. In other words, the tVNS-induced increase in LC-NE activity appeared effective only when it occurred alongside a difficulty to decide accurately. Notably, this aligns with the idea that a key principle guiding LC-NE function is the optimization of energy expenditure, with LC selectively supporting processes essential to task goals while ignoring, or even inhibiting those that have minimal impact on task performance [27, 28]. Consistently, past studies have reported a specific contribution of LC in demanding contexts [11-13, 82, 83], such as during the exertion of a physical [11, 83-85] or cognitive effort [14, 24]. Therefore, it is likely that tVNS in our study specifically enhanced decision-making processes when the task was more demanding, resulting in a selective effect in those participants for whom performing after errors was particularly challenging.

The application of tVNS in our study induced a systematic and consistent increase in pupil dilation, indicating a reliable boost in LC-NE activity across all tVNS trials of the RDM task. Therefore, the selective effect of tVNS on accuracy cannot be attributed to varying tVNS impacts on LC-NE activity. Instead, it appears that, depending on task demands, the tVNS-induced boost in LC-NE activity produced differential effects on the regions involved in the decision-making processes. This is further supported by the variation we found in the drift rate parameter of the DDM, which reflects the efficiency of evidence accumulation in visual areas, and which closely paralleled the accuracy changes observed at the behavioral level. Specifically, in less proficient participants, the drift rate declined after errors in SHAM blocks, indicating impaired evidence accumulation. However, tVNS appeared to prevent this decline, as no reduction in drift rate was observed following errors in tVNS blocks. In contrast, more proficient participants showed stable drift rates after errors in SHAM blocks, with tVNS having no impact on this parameter. This suggests that the tVNS-induced LC-NE drive selectively enhanced visual processing to counteract post-error impairments in evidence accumulation. The fact that tVNS had no effect on the non-decision time parameter in the DDM suggests that it may not have influenced the motor processes underlying decision-making. To clarify this, future studies should investigate decision-making using responses that involve more ecologically valid movements, such as reaching towards a left or right target, rather than relying on a simple keypress.

We also observed intriguing effects on the decision boundary parameters of the DDM. In SHAM blocks, the boundary intercept increased (as a trend) after errors, reflecting a higher, more conservative threshold for decision-making, consistent with previous research showing that participants accumulate more evidence before making decisions after a mistake [45]. However, this shift was mitigated to some extent because the boundary slope became steeper after errors, indicating a faster collapse of the decision threshold as the trial progressed [94]. Despite these opposing adjustments, the net effect was a decrease in decision speed, as evidenced by generally longer RTs after errors compared to correct decisions. This aligns with the post-error slowing literature [87, 88], indicating a shift in decision-making dynamics after errors. However, this shift did not reflect increased caution here, as accuracy did not improve after errors: in fact, it even declined among less proficient participants. Now, what about the impact of tVNS on the boundary parameters? Based on the lack of tVNS effect on RTs, we predicted that we would not observe any change in boundary parameters. But surprisingly, both the boundary intercept (as a trend) and slope were reduced following errors compared to SHAM blocks, suggesting that tVNS lessened the post-error adjustments in decision boundaries observed in the SHAM blocks, even though RTs did not change. A plausible explanation for the absence of RT change is that the effect of tVNS on the intercept and slope was perfectly balanced, leading to no net change at the behavioral level. Nevertheless, we acknowledge that tVNS seems to have influenced parameters typically linked to urgency in decision-making. Future studies should explore this further, perhaps using tasks that emphasize response speed. We believe that if RTs become a critical factor in task performance (where accuracy was the main focus here), the tVNS-induced boost in LC-NE activity might reveal additional benefits at that level too.

## Conclusion

Our study demonstrates that tVNS is effective in directly stimulating the LC-NE system, offering a promising new approach to investigate the role of this neuromodulatory system in human behavior. This technique goes beyond the limitations of correlational observations based on pupil size [9, 14]. Moreover, probing the system causally appears particularly important now that we observed a dissociation between the effects of tVNS on pupil dilation and the specific behavioral outcomes: behavioral regulations were not consistently accompanied by variations in pupil size. Additionally, our findings reveal that tVNS enhanced accuracy without affecting RTs, which supports the “gain” hypothesis over the “urgency” hypothesis.

Notably, the improvement in accuracy was limited to more demanding task conditions, suggesting that the LC-NE system adjusts its contribution based on task complexity, as previously proposed. However, this also leaves open the possibility that the LC-NE system may still assist processes generating urgency when this aspect of behavior becomes particularly critical, a question for future research. Overall, this study marks an important step in leveraging tVNS as a non-invasive tool to explore the causal role of the LC-NE system in human behavior.

