## Supplementary Material for "Transcutaneous Vagus Nerve Stimulation Boosts Post-Error Accuracy During Perceptual Decision-Making"

### Supplementary Materials

**Experimental procedures**

***Experiment 1***

Participants focused on a white cross in the center of a black screen while experiencing tVNS/SHAM stimulation at rest. Each participant completed 4 to 6 blocks of 20 trials with either tVNS or SHAM stimulation in separate blocks, providing at least 40 trials per condition, which is sufficient for reliable pupil measurements [1]. Stimulation periods were separated by a 10-11 second interval, making each trial lasting 14-15 seconds. The order of tVNS and SHAM blocks was alternated and counterbalanced across participants.

***Experiment 2***

Each experiment started with a training session, where participants completed 4 blocks of 40 trials, with coherence decreasing progressively (c’ = 0.8, 0.4, 0.2, and 0.1, respectively). Subsequently, a dot coherence calibration session began, aiming to determine the individual c’ value corresponding to 70% of accuracy. During this session, participants achieved 2 blocks of 100 trials, comprising randomly interleaved stimuli with varying coherence levels (c’ = 0.1, 0.2, 0.4, 0.6 and 0.8, 40 trials each). Following completion, each individual’s psychometric function was estimated using the proportional-rate diffusion model [2]. The c’ value corresponding to 70% accuracy was interpolated from this function and utilized in the main experiment blocks. Afterwards, each subject underwent a second calibration session to select their individualized intensity for tVNS and SHAM stimulation.

During the main experiment blocks, subjects performed the RDM task (implemented in Matlab R2023a using the Psychophysics toolbox [2]) across 8 blocks of 40 trials, split into 4 blocks of tVNS and 4 blocks of SHAM stimulation. The tVNS and SHAM blocks were alternated, with the sequence of the first 4 blocks reversed in the second set of 4 blocks. These sequences were counterbalanced across subjects. Subjects were free to rest between the blocks.

**Pupil data pre-processing**

Blink periods were detected and removed through linear interpolation of values 100 ms before and after each identified blink, as implemented in custom-made Matlab scripts. The pupil data were then low-pass filtered using a 10Hz fourth-order Butterworth filter with zero-phase shift. Trials where interpolated data accounted for more than 50% of data points were excluded.

**Drift diffusion model (DDM) fitting and parameter recovery**

The DDM was fitted to the RT and Accuracy data using a variant that includes linearly collapsing decision boundaries. The decision boundary during each trial was modeled as:

$$a_{t}=a_{0}-ut$$

where $a_{t}$ is the boundary height at time $t$, $a_{0}$ is the initial decision boundary (further referred to as boundary intercept), and $u\geq0$ is the collapse rate of the boundary (further referred to as boundary slope). A lower boundary intercept can be interpreted as a higher initial urgency while a steeper boundary slope can be seen as a stronger increase in urgency to respond as time progresses, leading to a less skewed RT distribution and a drop in accuracy over time [3]. The drift rate, boundary intercept, boundary slope, and non-decision time were estimated for each participant based on Block-Type (tVNS, SHAM) and Trial-History (after-correct, after-error).

Quantile optimization was used for fitting the DDM [4]. The optimization was performed by a differential evolution algorithm as implemented in the DEoptim package in R [5]. The population size was set at ten times the number of free parameters. Convergence was met if no improvement in the objective function was found in the last 100 generations. To ensure a reliable interpretation of estimated parameters, a parameter recovery analysis was performed. We simulated the data of 1000 simulated participants using parameter values randomly sampled from their typical range. We managed to recover the parameters with as few as 25 trials: all correlations between generated and estimated parameters rs > 0.73, except for the boundary slope (r = 0.53). As a result, we excluded 6 subjects who had less than 25 trials after errors, leading to 36 ± 10.4 and 40 ± 12.1 remaining trials after errors in tVNS and SHAM conditions of 15 subjects, respectively.

**Pearson’s partial correlation analysis on behaviors**

To further explore the relationship between tVNS effects and the accuracy decline following errors, we conducted a Pearson’s partial correlation analysis. This analysis evaluated the relationship between the tVNS effect on post-error accuracy (delta Block-Type: tVNS minus SHAM) and the accuracy drop after errors in SHAM blocks (delta Trial-History: after-error minus after-correct). The results revealed a significant negative correlation (R = -0.536, p = 0.015, Supplementary Fig. 1A), indicating that participants who showed a larger accuracy decline after errors in SHAM blocks benefited more from tVNS in terms of accuracy improvement. Similarly, we run the same correlation analysis on RT. As expected from the absence of Block-Type effect, we did not find any Pearson’s partial correlation (R = -0.262, p = 0.264, Supplementary Fig. 1B) between the degree of post-error slowing in SHAM blocks (delta Trial-History: after-error minus after-correct) and a potential tVNS effect on RT after errors (delta Block-Type: tVNS minus SHAM).

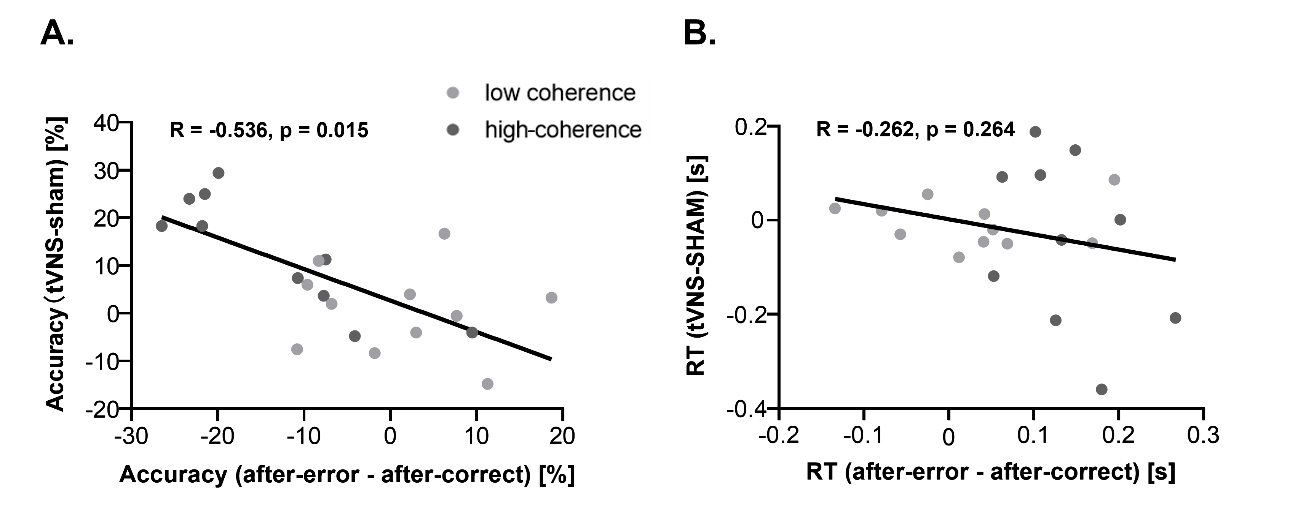

**Supplementary Figure 1. Pearson’s partial correlation analysis on behaviors.** Data in low-coherence and high-coherence subjects were displayed with light and dark gray dots, respectively. **A. Correlation on Accuracy.** The correlation between the tVNS effect on accuracy after errors and the accuracy drop following errors in SHAM blocks: subjects who experienced a greater accuracy drop after errors also benefited more from tVNS in terms of accuracy improvement. **B. Correlation on RT.** No correlation between the tVNS effect on RT after errors and the post-error slowing in SHAM blocks.

**Delta values of drift-diffusion model (DDM) parameters**

To further investigate post-error adjustments and tVNS effects, we computed Delta values (after-error minus after-correct trials) for key DDM parameters: drift rate, boundary intercept, boundary slope, and non-decision time between tVNS and SHAM conditions.

*Drift Rate:* The high-coherence group displayed a lower (negative) Delta drift rate compared to the low-coherence group (t(28) = 3.31, $p_{holm-bonferroni}$ $p_{holm-bonferroni}$ = 0.012, Supplementary Fig. 2A). But then subjects in the former group displayed a normalization of their Delta drift rate (back to values close to 0) in tVNS blocks (t(14) = 2.97, $p_{holm-bonferroni}$ $p_{holm-bonferroni}$ = 0.030), an effect absent in the low-coherence group (F(1, 21) = 0.11, p = 0.740).

*Boundary intercept:* Deltas of boundary intercept were higher in SHAM blocks than in tVNS blocks (F(1, 15) = 7.43, p = 0.02, Supplementary Fig. 2B), indicating the absence of post-error increase in boundary intercept with tVNS. Note that these observations were made regardless of Coherence-Group.

*Boundary slope:* Deltas of boundary slope were higher in SHAM blocks than in tVNS blocks (F(1, 15) = 18.10, p < 0.001, Supplementary Fig. 2C), indicating the absence of post-error increase in boundary slope with tVNS. Here too, these observations were made regardless of Coherence-Group.

*Non-decision time:* We did not find the Block-Type effect on Delta non-decision time (F(1, 30) = 2.74, p = 0.108, Supplementary Fig. 2D).

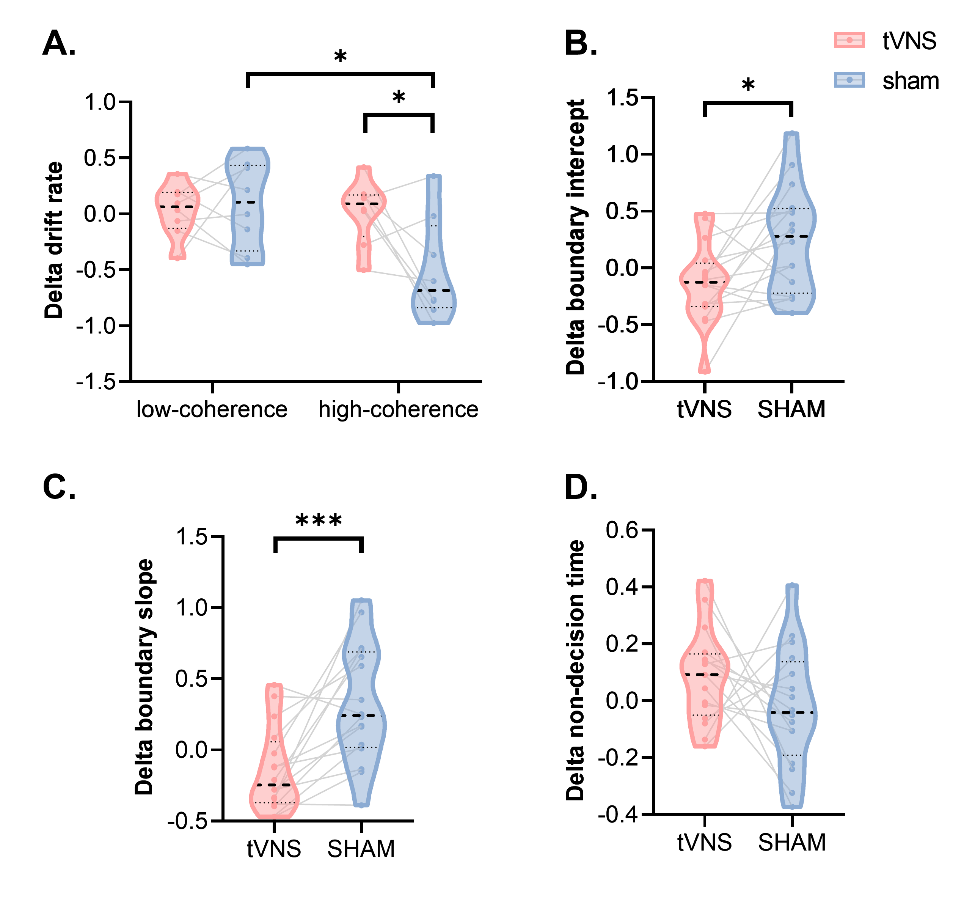

**Supplementary Figure 2. tVNS effect on Delta Values of Drift Diffusion Model (DDM) Parameters.** The graphs display Delta values of DDM data, separated for tVNS (red) and SHAM (blue) blocks, with median values featured as black dotted lines, distributions as violin plots, and individual data as thin gray lines. **A. Delta drift rate.** For the high-coherence group selectively, note the lower Deltas in SHAM blocks (reflecting an after-error drop in drift rate), which then returns close to 0 with tVNS. **B. Delta boundary intercept.** Note the higher Deltas in SHAM blocks (reflecting an after-error increase in boundary intercept) than in tVNS blocks. **C. Delta boundary slope.** Note the higher Deltas in SHAM blocks (reflecting an after-error steepening of the boundary slope) than in tVNS blocks. **D. Delta non-decision time.** Note the consistent Deltas across conditions. *: $p$ < 0.05. ***: $p$ < 0.001.
